# Fine-Tuning of Label-Free Single-Cell Proteomics Workflows

**DOI:** 10.1101/2025.11.17.688849

**Authors:** Pauline Perdu-Alloy, Keller Charline, Anjali Seth, Christine Carapito

## Abstract

Mass spectrometry-based single-cell proteomics emerges as the most promising method for studying cellular heterogeneity at the global proteome level with unprecedented depth and coverage. Its widespread application remains limited due to robustness, reproducibility, and throughput requirements, still difficult to meet as analyzing large cohorts of single cells is necessary to ensure statistical confidence. In this context, we have conducted method optimizations at three levels. First, we benchmarked three distinct workflows compatible with the nanoElute2 platform using different sample collection/preparation plate supports (EVO96 oil-free, LF48 oil-based and LF48 oil-free, a streamlined automated sample resuspension and direct injection protocol). Then, we compared the optimized EVO96 workflow on nanoElute2 with EVOSEP-based separations operating at two analytical throughputs (80 and 120 samples per day). Subsequently, we evaluated digestion efficiency using a range of enzyme/protein ratios (1:1; 10:1; 20:1; 50:1) to maximize peptide recovery. Finally, the chromatographic setup was refined to determine the best compromise between throughput and robustness. Altogether, these optimizations allowed to establish a robust workflow quantifying up to 5,000 proteins in 10min gradient time per single Hela cell at a 55 samples-per-day throughput.

## INTRODUCTION

Qualitative and quantitative analysis of the whole proteome highlight the response of an organ or population of cells to disease or treatments. However, under varying conditions and overtime, multicellular organisms exhibit great heterogeneity in their protein expression patterns^1,2,3^. In this complexity, bulk protein analysis is unable to distinguish healthy from diseased/infected cells^4,5,6^ nor to identify subclonal populations that behave differently from the rest of the cell population (e. g. in cancer)^7,8,9,10^. Thanks to constant technological progress, investigation of single cells’ proteome now becomes feasible, but still faces analytical challenges, mainly due to its need for sample miniaturization (picoliter volume dispense), highly sensitive instrumentation (enabling detection of 80-400pg protein input material), and sophisticated computational tools (for decomplexification of high throughput datasets ^11,12^).

Tailored sample preparation protocols are a crucial step of the workflow and since its introduction in 2017, the cellenONE robot has transformed single-cell sorting using an image-based technology to select cells based on their morphological parameters, without the need for labeling. Combined with recent breakthroughs in ion mobility-mass spectrometry using ultra-fast and ultra-sensitive mass analyzers, latest generation couplings have allowed significant improvement of the resolution and depth of single cell proteome analysis. In particular, the creation of new precursor scan designs and their fragmentation management in tandem with their accumulation, enabled by the DIA-PASEF (Data-Independent Acquisition-Parallel Accumulation Serial Fragmentation) strategy^13,14,15^ enhance selectivity and efficiency.

In this context, we have undertaken a series of method optimizations at all steps of the workflow to further increase single cell proteome depth and coverage.

## EXPERIMENTAL PROCEDURES

### Single cells preparation on the cellenONE system

In house cultured HeLa cells were resuspended to a final concentration of approximately 200cells/µL in DPBS, and then loaded into a piezoelectric dispensing M-size capillary in the cellenONE system (Cellenion, Lyon France), with droplet generation settings of 97 V, 48µs and 500 Hz.

A 300 nL solution of Mastermix^16^ containing lysis and digestion reagents (100 mM TEAB, ref: 15658320 (ThermoFisher Scientific); 0.2% DDM, 850520P-1G (Sigma-Aldrich); 10 ng/µL trypsinLysC, ref: V5072 (Progema Corporation) was dispensed into each well of the substrate of choice (either an EVO96, an LF-48 with oil or an oil-free LF-48 proteoCHIP) at 10°C with 50% humidity. Single-cell sorting followed fixed parameters for optimal selection: intensity min 1 and max 255; elongation max 2; diameter min 16µm and max 40µm. The detection was set for diameter min 2µm and max 40µm and for elongation max 4. This isolation step was performed at 10°C with 50% humidity.

After dispensing, single cells were incubated at 50°C for 1.5 hours from 10°C to 20°C at 75% humidity and from 20°C to 50°C at 85% humidity, with rehydration cycles (260 nL/well, 500 Hz) to prevent evaporation of the lysis buffer. Samples were then cooled to 20°C with 85% humidity, maintaining rehydration cycles throughout. Digestion was then halted by manually adding 3.2 µL of 2% acetonitrile (ACN) and 0.1% formic acid (FA) per well at room temperature, except for LF48 oil-free for direct injection, as the nanoElute2 system automatically performs the dilution step.

For EVO96 support injected on the nanoElute system, thanks to a centrifugation step at 700g for 30sec, the peptide mixture was then transferred and collected to a 96-well injection plate, which was stored at −80°C before mass spectrometry analysis. For EVO96 support injected on the Evosep system, after EVOTIPS clean-up procedure, peptides were loaded on the EVOTIPS and cleaned-up by centrifugation at 800g for 60sec. The peptide mixture was then analyzed by mass spectrometry. For LF48 oil-based support, after a cooling step at 4°C in a fridge, the liquid peptide mixture is collected and transferred to a 96-well injection plate, which was stored at −80°C before mass spectrometry analysis. For LF48 oil-free support, the plate is collected directly after the incubation step and ready for injection.

### Nano-liquid chromatography

Nanoelute system. Peptides mixtures were separated on a 5cm C18-RP analytical column (75µm inner diameter, 1.7µm particle size, AuroraRapid, IonOptiks) heated up to 50°C using the nanoElute2 nanoliquid chromatography system (Bruker Daltonics, Bremen, Germany). Separation was performed using a linear gradient from 5 to 35% of solvent B (solvent A: 0.1% FA in H_2_O and solvent B: 0.1% AF in ACN) over a 10min gradient at 0.25µL/min flow rate. Column washing and regeneration were performed in 6 minutes by ramping the percentage of mobile phase B from 35% to 90% and then back to 5%.

Evosep system. Peptides mixtures were separated with the same column as for the Nanoelute system on the EVOSEP One system (Evosep Biosystems, Odense, Denmark). Peptides separation was performed using Whisper Zoom methods 80 SPD and 120 SPD. Separation was performed using a gradient from 0 to 40% of solvent B (solvent A: 0.1% FA in H_2_O and solvent B: 0.1% AF in ACN) respectively over a 16.3 min and a 10.3 min gradient at 0.2 µL/min flow rate.

### Tandem mass spectrometry

Separated peptides were analyzed on a timsTOF Ultra 2 mass spectrometer (Bruker Daltonics, Bremen, Germany) operated in diaPASEF mode. The instrument settings included a capillary voltage of 4.5 kV, with a gas flow rate maintained at 3.0 L/min at 200 °C. Fragmentation windows, with a width of 25 m/z, were established from 0.64 to 1.37 V.s/cm^2^ along the ion mobility range, covering a mass-to-charge ratio (m/z) range from 400 to 1000. The accumulation time for each ion mobility scan was set at 100 ms, with a ramp time of 100 ms. Collision energies were applied across the ion mobility range, starting from 0.60 V·s/cm^2^ at 20 eV and increasing to 1.60 V·s/cm^2^ at 59 eV to optimize ion fragmentation. Each cycle, lasting 0.96 seconds, comprised a full MS1 scan, followed by 24 MS/MS windows. Within each of these windows, ions were fragmented across 8 MS/MS ramps. Complete dataset has been deposited in the ProteomeXchange Consortium via the PRIDE partner^17^ repository with the dataset identifiers PXD067019.

### Data treatment

Raw instrument files collected from DIA-PASEF were analyzed through DIA-NN^18^ (v. 1.8.2beta27). The processing was performed using a library-free mode. *In silico* digestion using Trypsin/LysC was performed on a human proteome fasta (20,409 reviewed Swiss-Prot entries) allowing a maximum of 1 missed cleavage. Theoretical peptide spectra were predicted according to their retention time and ion mobility thanks to a deep learning-based algorithm provided by DIA-NN ^19^. For peptide spectrum matches (PSMs), precursors were filtered at 1% FDR. Variable modifications on peptides were set to methionine oxidation and N-term acetylation with a maximum of 5 variable modifications. For data processing, the peptide length ranged from 7-30 amino acids; precursor charge ranged from 1-4 whereas their m/z ranged from 300-1800 and finally, fragment ions m/z ranged from 200-1800. All mass accuracies, notably the precursor masses (MS1) and fragment masses (MS2), were defined at a tolerance of 10ppm. Match between run (MBR) was checked for cross-run analysis considering single cell replicates only.

## RESULTS AND DISCUSSION

### 1. Sample preparation plate support optimization

Single cell proteome preparation workflows consist of five main steps: (1) lysis and digestion mixture (mastermix) dispensing, (2) single-cell isolation, (3) incubation, (4) peptide dilution, and (5) most often peptide transfer into an injection plate. Steps 1 to 3 are common to every workflow. In our case, the mastermix contains Trypsin-LysC at 10ng/µL, 0.2% n-dodecyl β-D-maltoside (DDM), and 100mM triethylammonium bicarbonate (TEAB) in initial concentrations. Single cells are isolated into the collection plate wells, followed by an incubation step of 1.5 hours at 50°C, allowing for cell lysis, protein extraction, and digestion into peptides.

To assess potential peptide losses and sources of human-introduced variability –primarily caused by evaporation, solvent dilution (i. e., a small final peptide volume of 0.3µL diluted in a relatively large recovery solvent volume of 3.2µL), and manual pipetting– four workflows based on various sample collection plate supports were benchmarked as illustrated in **Figure 1**:

**Figure 1.**
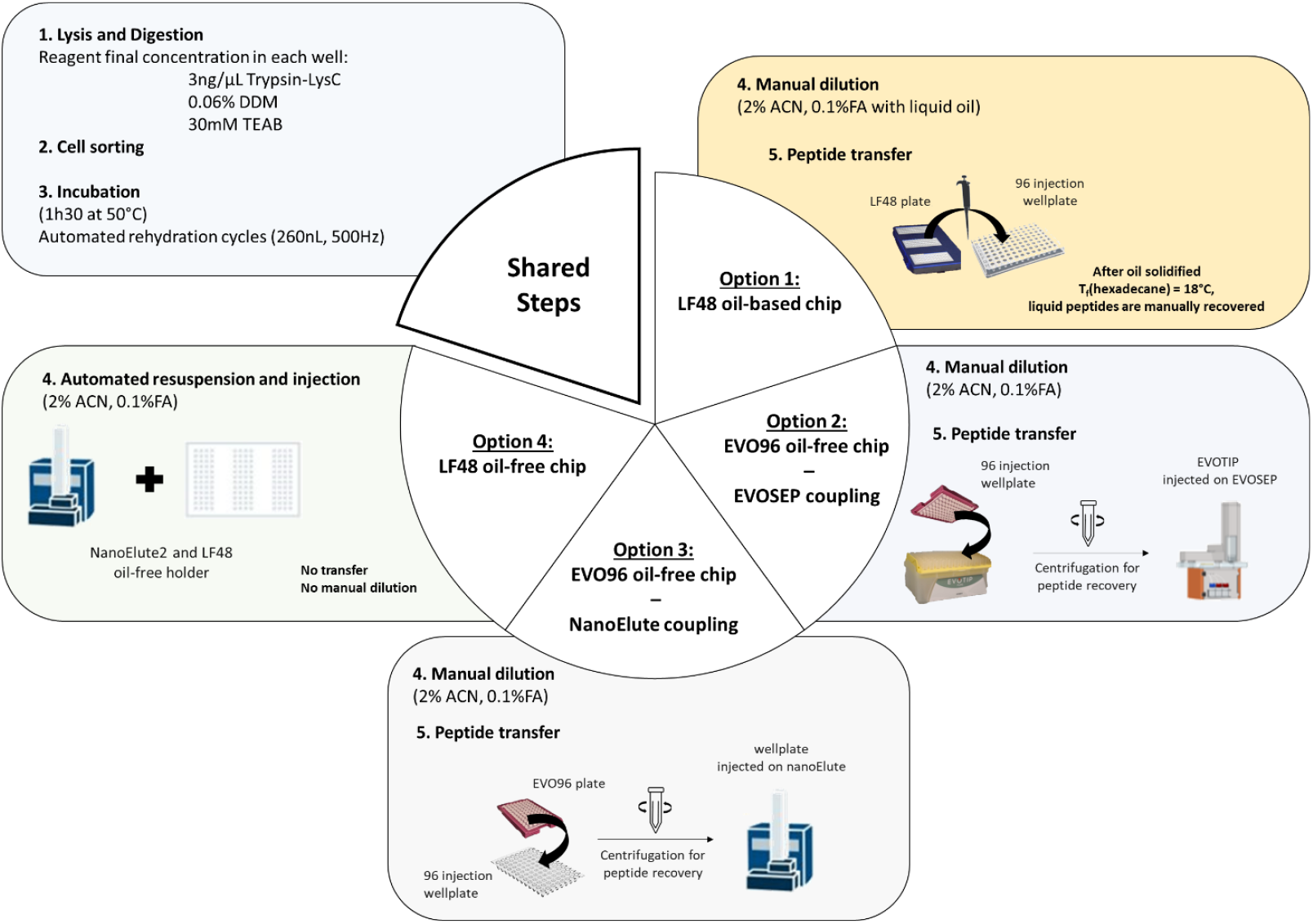
Four single cell sample preparation protocols benchmarked using the cellenONE sorter/liquid dispenser. Lysis and digestion, followed by incubation with automated rehydration cycles are common steps to all three workflows. Then, option 1 relies on a LF48 Oil-Based plate support involving a manual dilution step in liquid oil, followed by a manual peptide transfer after oil solidification (Tf (hexadecane) = 18°C); option 2 uses an EVO96 Oil-Free plate support, involving manual dilution followed by peptide loading onto pre-conditioned Evotips after centrifugation for injection on the EVOSEP system; option 3 employs the same EVO96 oil-free support, repurposed for nanoELute injection, where peptides are transferred into a 96-well injection plate via centrifugation; option 4 relies on a LF48 Oil-Free plate support suited for automated resuspension and direct injection on a nanoElute 2 system.

- **Option 1** – The LF48 oil-based support, the original cellenONE protocol, for the collection of 48 single cells per chip, integrating a thin hexadecane layer requiring a final manual sample transfer.
- **Options 2 and 3** – The EVO96 support, an oil-free proteoCHIP for the single-cell collection (of up to 96), with hands-free peptide transfer via a short centrifugation step. Although originally developed for coupling with the EVOSEP system (Option 2), we also attempted its adaptation for nanoElute injections (Option 3, EVO96_nE).
- **Option 4** – The LF48 oil-free support enabling automated resuspension and injection, using a custom-designed proteoCHIP holder compatible with the nanoElute2 autosampler.

Figure 2 summarizes the results obtained with the three evaluated workflows compatible with the nanoElute 2 platform (Options 1, 3, and 4). In terms of proteome coverage, among the three evaluated workflows, the LF48 oil-free direct injection protocol enables the highest coverage with a median and average 4,682 / 4,437 protein groups and 30,995 / 29,411 peptides identified.

The lowest protein groups/peptides (2,114 / 2,064; 10,581 / 10,792 median and average) numbers are reached with the EVO96 collection plate support, while the LF48 oil-based protocol shows intermediate coverage performances (3,114 / 3,109; 17,774 / 18,065 median and average protein groups/peptides, respectively) (**Figure 2A, B**). Importantly, although MBR increases the number of identifications by reducing the number of missing values with elution profile recovery from other single cells analyzed in parallel, the relative performance trend between workflows remained unchanged. It confirms that the differences observed are robust and not partially due to the MBR processing. Furthermore, the EVO96 workflow offers an efficient and user-friendly format thanks to the centrifugation step, which eliminates manual pipetting and reduces hands-on time allowing for a total preparation time of ~3 hours for 192 cells (96×2plates) but the trade-off is a slight reduction in sensitivity when compared to the LF48 plate supports.

**Figure 2.**
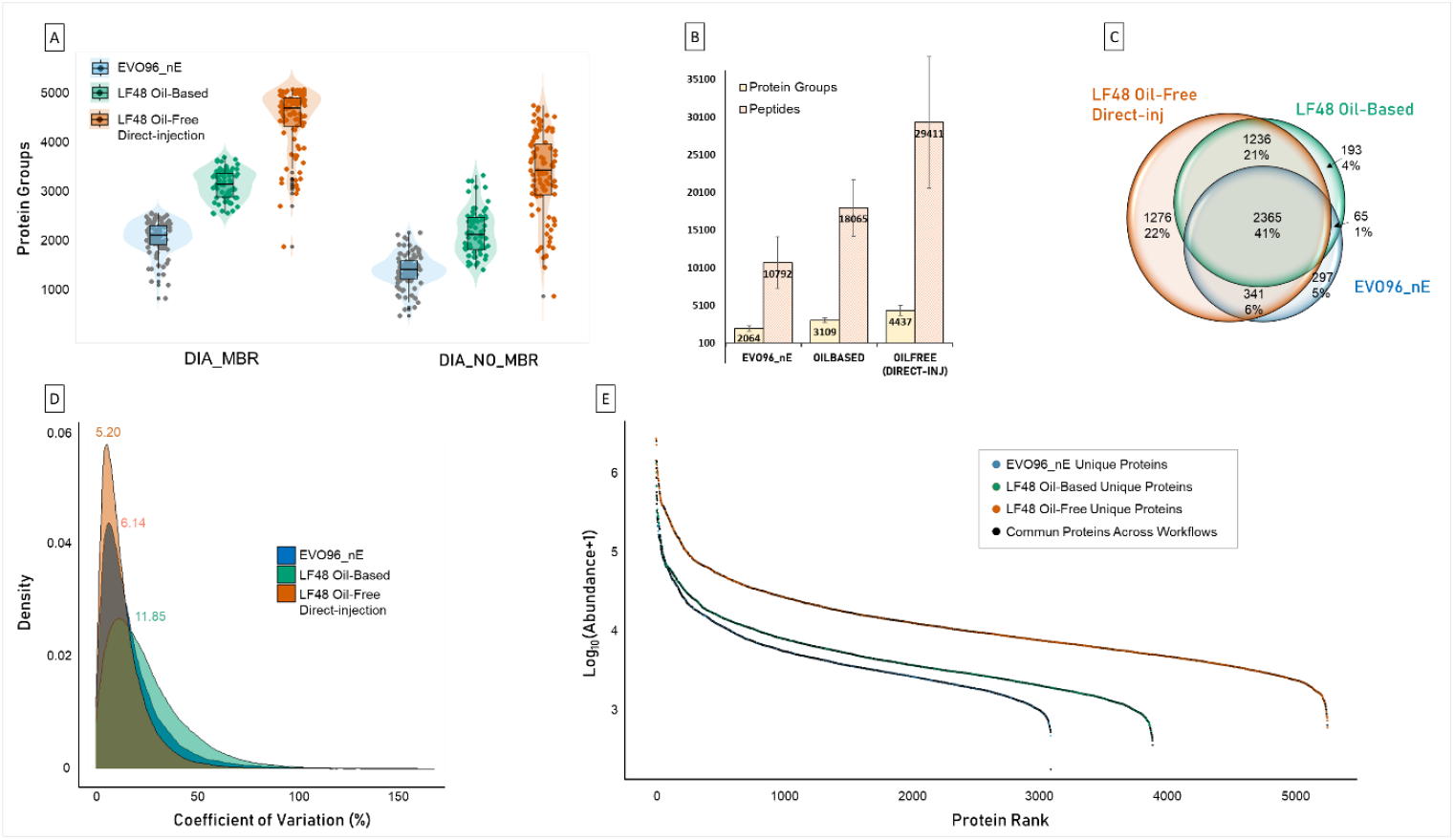
Benchmarking of three single-cell proteomics workflows (EVO96 (blue), LF48 oil-based (green), and LF48 oil-free (orange)) run on nanoElute 2 (A) Violin plots representing the number of protein groups identified across single cells for each tested workflow using DIA-NN with MBR (left) and without MBR (right). (B) Protein groups and peptide number identified across the three benchmarked workflows. (C) Venn diagram showing proteins’ overlap across the three workflows. (D) Coefficients of variation (CVs) density distributions across the three sample preparation workflows. (E) Protein abundance ranking plot comparing the dynamic range of protein groups across the three workflows. Black dots correspond to the shared proteins across the condition whereas the colored ones correspond to unique proteins.

The higher coverage achieved with the oil-based LF48 plate support reflects the benefit of hexadecane oil layer, which, in addition to the regular and automatic water dispensing aimed at minimizing evaporation, provides an additional protective layer reducing evaporation effects. However, the highest coverage is reached with the oil-free direct injection LF48 support which highlights the strong negative effects of sample dilution and sample transfer (whether done manually or by centrifugation). Indeed, thanks to the adapter plate support compatible with the nanoElute2 automater, single cell protein extracts can be directly resolubilized and injected from the well, without any intermediate transfer or dilution step. This streamlined workflow minimizes sample handling and eliminates peptide losses associated with adsorption to surfaces, evaporation, or pipetting. However, a noticeable drawback of this workflow is that it shows more outliers than the two other protocols, possibly attributable to time-dependent effects (**Suppl. Figure 1**) –suggesting that further optimizations— particularly in plate preservation and injection throughput—could help limit this drawback.

In terms of proteome overlap, the oil-free LF48 resulted in the highest proportion of unique protein identifications (22%), while 2,365 protein groups (41%) were shared among all three conditions (**Figure 2C**). Proteins uniquely identified with the oil-based LF48 and EVO96 are marginal and represent less than 5% of the total proteins.

In terms of quantitative reproducibility, the coefficients of variation (CV) density distributions revealed lower variability in the LF48 oil-free condition, followed by the EVO96 workflow (**Figure 2D**). The LF48 oil-based workflow showed the highest variability, suggesting that despite its evaporation-protective role, the manual peptide transfer introduces additional and unavoidable variability. Oil-free and EVO96 plate supports show good and equivalent performances in terms of quantification reproducibility. The LF48 oil-free condition also achieved the widest dynamic range, enabling the detection of lower abundant proteins (**Figure 2E**).

To further assess the nature of identified peptides, hydrophobicity of each detected peptide sequences was evaluated using the GRAVY (Grand Average of Hydropathy) score across the three workflows. The GRAVY score is calculated as the average of the hydrophobicity values of all amino acids in a peptide sequence according to the Kyte and Doolittle scale^20^. Thus, a positive GRAVY score indicates the presence of more hydrophobic peptides; while a negative score indicates the presence of more hydrophilic peptides. **Suppl. Figure 2** shows the GRAVY score average for each single cell across the three workflows. Although the resulting scores are within the same range (from −0.6 to 0.25), it is interesting to note that the EVO96_nE workflow tends to lead to more hydrophobic peptides recovery (with an approximate median GRAVY of −0.1), while the LF48 workflows cover a higher proportion of hydrophilic peptides (with an approximate median GRAVY of −0.4). These differences may reflect the composition of the injection plates: as the Teflon plates used in LF48 could inherently favor the loss of hydrophobic peptides, while the polypropylene plates used in EVO96 could favor the loss of hydrophilic ones.

Moreover, we also considered the number of peptides carrying methionine oxidations (UniMod:35). The distribution of oxidized peptides differs depending on the workflow (**Suppl. Figure 3**). Indeed, the EVO96_nE and LF48 direct injection workflows show lowest median numbers of oxidized peptides (80±39; 35±12 respectively), in contrast to the LF48 oil-based workflow (using hexadecane) that shows a wider distribution and greater heterogeneity (119 ± 80) concerning this oxidation phenomena. This suggests that the presence of oil during the preparation process may increase the variability of methionine oxidations between single-cells, potentially underlying differences in peptide exposure to oxidative conditions.

**Figure 3.**
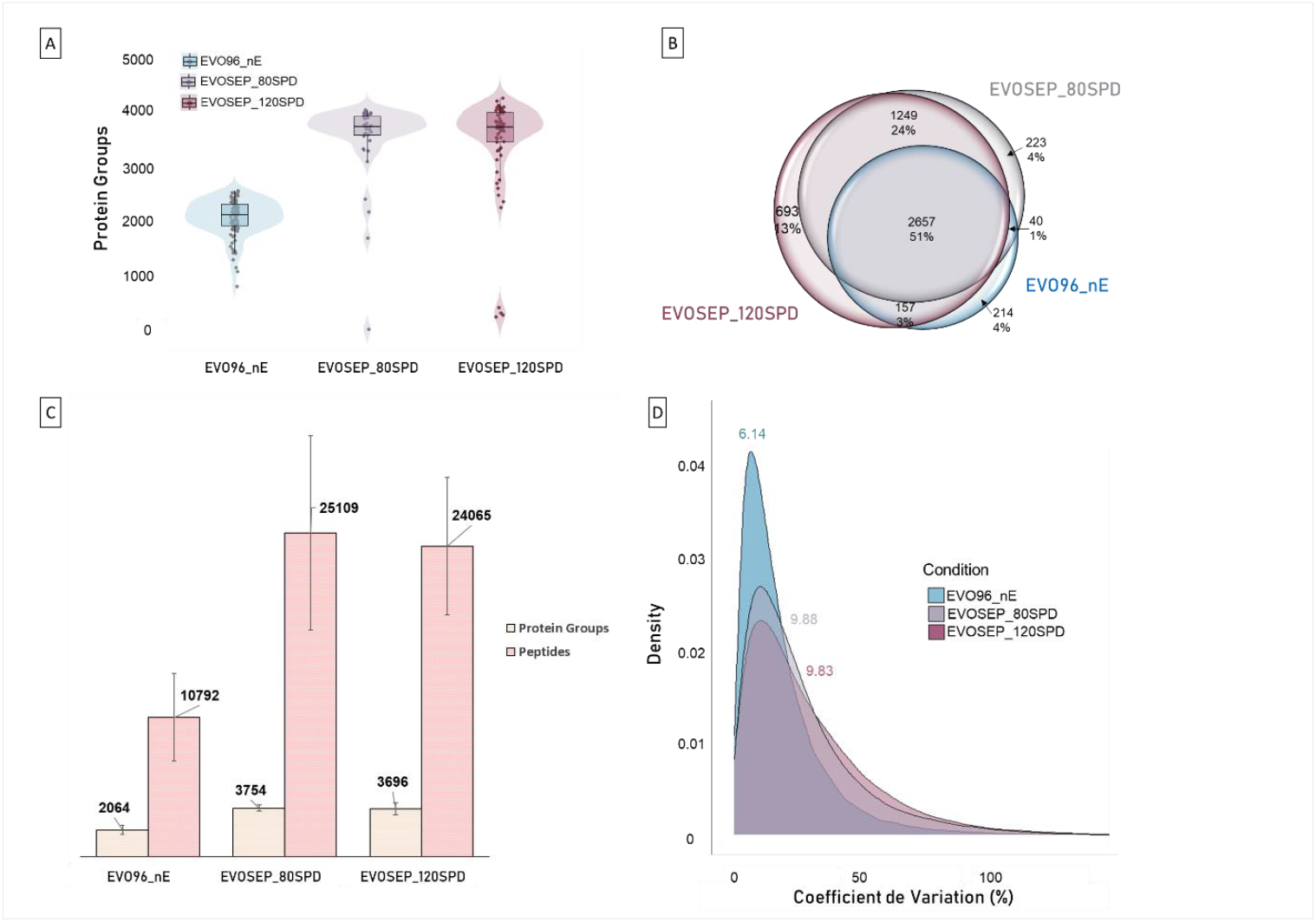
Comparison of SCP EVO96-based workflows across two chromatographic systems. EVO96_nE was analyzed on a nanoElute 2 system (N=96 single-cells) compared to two EVOSEP workflows that were acquired using two sample throughputs (80 and 120SPD with N=40 single cells each). (A) Violin plots representing the number of protein groups identified per single cell for each workflow (EVO96_nE in blue, EVOSEP80SPD in grey, and EVOSEP120SPD in purple). (B) Venn diagram showing the overlap of identified protein groups across the workflows. (C) Bar plots giving the total numbers of identified protein groups (orange) and peptides (pink) for each workflow. (D) Density distributions of protein abundance coefficients of variation (CVs) across the three workflows.

As a conclusion, we demonstrate hereby that every single step of the sample preparation workflow can drastically alter identification results and drives to a significant loss of proteome coverage and robustness. Sample preparation needs to be finely tuned. Furthermore, we are aware that the use of the EVO96 well plate coupled with nanoElute is diverted from its original purpose, which is generally to be integrated into a downstream cleanup analysis on the EVOSEP system. Therefore, we compared the EVO96 workflow on the two chromatographic systems in the following section.

### 2. Comparison of the EVO96 workflow on two chromatographic platforms

Then, we compared the EVO96 plate adapted to nanoElute with the original EVOSEP-based workflow at two different analysis throughputs, specifically 80 and 120 samples per day (SPD). To ensure a fair comparison, the EVO96_nE workflow was performed at a “80-SPD-like” throughput, which closely matches the analysis gradient of the 80SPD on EVOSEP.

Our results, presented in **Figure 3**, show a significant increase in the number of protein and peptide identifications across all EVOSEP workflows. In contrast to the EVO96_nE workflow that delivers 2,064 protein groups and 10,792 peptides; the EVOSEP workflow identified 3,754 / 3,696 and 25,109 / 24,065 protein groups / peptides at 80SPD and 120SPD, respectively.

Significant increase of approximately 80% in protein identifications and of up to 130% at the peptide level was observed using EVOSEP when compared to the EVO96_nE workflow. This increase confirms optimized design of the EVO96 plate for the EVOSEP system. Its tips, EVOTIPs, work as a SPE-like purification and preconcentration step, protecting the nanoLC system and ensuring that, unlike nanoElute injections, where the entire cell content is loaded on column in direct injection mode, only the eluted peptide fraction is predominantly transferred to the mass spectrometer. This coupling allows reaching a high robustness and sensitivity with peptide recovery enhancement. In addition, thanks to this functioning, the EVOSEP platform does not require blank injections, while they are crucial on the nanoElute to avoid over-pressurization of the system by period washing. Thereby, analysis throughput can be increased.

### 3. Enzyme-to-Protein ratio optimization

We then focused on the lysis and digestion protocol, evaluating four enzyme-to-protein ratios (specifically 1:1, 10:1, 20:1 and 50:1). These ratios were intentionally chosen to not be in line with the lower trypsin concentrations commonly used in bulk proteomics (typically ranging from 1:100 to 1:10 (^21^). Given the need to work at the nanoliter scale, the lysis and digestion steps must be meticulously refined to accommodate for the extremely low protein amounts. Maintaining a high enzyme concentration is essential to ensure efficient digestion kinetics. In single-cell proteomics, protein digestion occurs within 1-2 hours, significantly faster than the overnight digestion commonly used in bulk workflows (^22,23,24^). In this context, optimizing digestion requires tuning of trypsin concentration, finding the right balance between minimizing trypsin autolysis, which can lead to signal suppression, and maximizing peptide generation to achieve better proteome coverage. While higher trypsin concentrations can enhance peptide generation, it also carries the risk of autolysis that could compromise the sensitivity of the analysis.

Both extreme ratios, namely 1:1 and 50:1, led to significant technical issues, notably numerous column blockages during chromatographic separation. Columns cloggings were attributed to either an excess of protease (in the 50:1 ratio) or an insufficient amount of protease (in the 1:1 ratio) to properly digest cellular proteins. Consequently, these two ratios were excluded from further analysis.

Results obtained for the other two evaluated enzyme-to-protein ratios, namely 10:1 and 20:1, are presented in **Figure 4**. Higher trypsin concentration resulted in higher proteome coverage with a 27% increase in peptide identifications (8,503 vs 10,792 peptides), a 10% increase in protein identifications (1,879 vs 2,064 protein groups) (**Figure 4A,B**), and a higher average number of peptides per protein (**Suppl. Figure 4**). One of the main challenges of single-cell proteomic studies is to ensure consistent sampling and quantification of thousands of proteins, while minimizing missing data^25^. Here, we observed not only fewer protein missed cleavages (1% less at 20:1) but also fewer overall missing values (36% at 10:1 compared with 32% at 20:1), indicating better digestion efficiency with more consistent quantification results (Fig S2).

**Figure 4.**
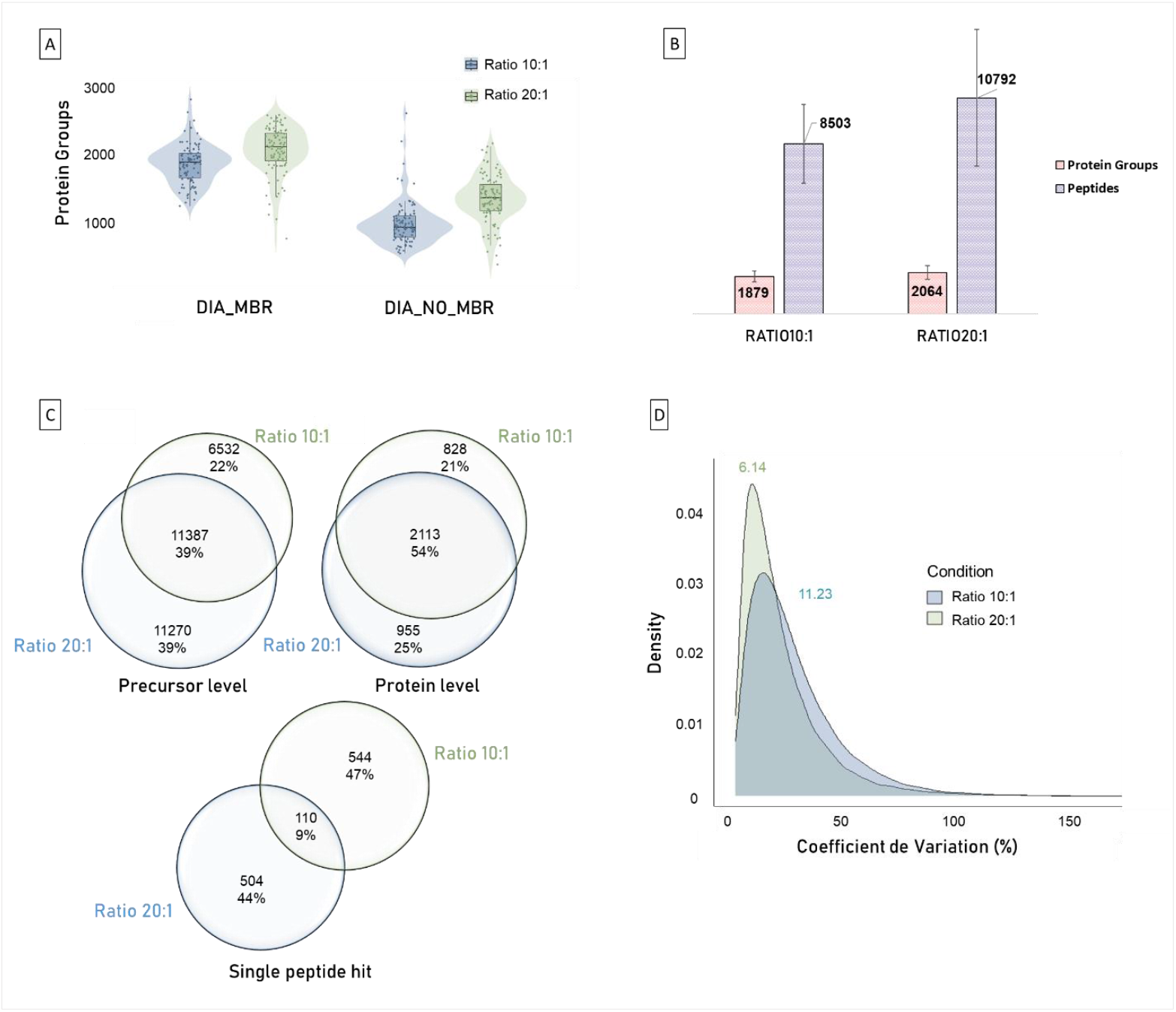
Evaluation of Trypsin-LysC digestion ratios. This figure presents a comprehensive analysis of HeLa single cells (N=96 cells per condition), comparing two digestion ratios (10:1 and 20:1). (A) Violin plots show the distribution of protein groups identified per cell under each digestion ratio, with or without Match Between Runs (MBR) processing. (B) Total number of identified protein groups and peptides at each of the two enzyme-to-protein ratio. (C) Venn diagrams illustrate the overlap in identified precursors (left), protein groups (right), and single-peptide hit proteins (down) between the two digestion ratios. (D) Density plots represent the distribution of coefficient of variation **(CV)** values across single cells.

**Figure 5.**
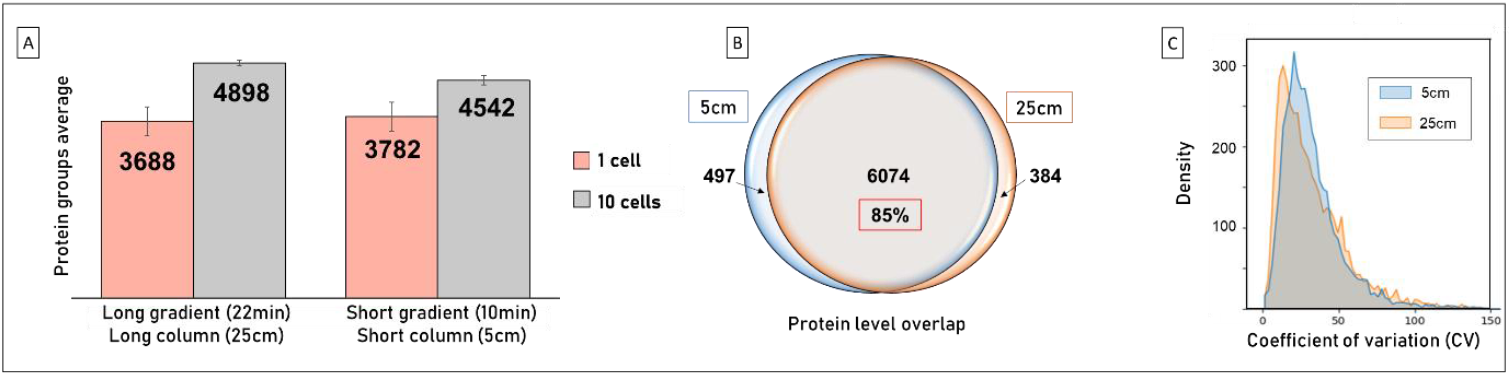
Gradient time and column length optimizations. (A) Number of protein groups identified from one single cell (pink) and ten cells (grey), analyzed using 5 cm and 25 cm IonOptiks separation columns. (B) Protein overlap between 5 cm (blue) and 25 cm (orange) separation column length for the single cells injected (N=10). (C) Density distribution of the CVs at the two tested conditions.

To assess digestion reproducibility, we also evaluated variability of the results across the hundred isolated cells per condition as a function of sample preparation condition. Distribution of the coefficients of variation (CVs) (**Figure 4D**) demonstrates that refining the workflow by adapting trypsin ratio is crucial for consistency of results. Indeed, the 20:1 ratio leads to a reduction in variability, as shown by the lower CV compared with the 10:1 ratio (maximum of CV density of 6.1 vs 11.2, respectively). In addition to this reduction in variability, critical for data interpretation, the 20:1 ratio presents also higher numbers of unique identifications at the protein and precursor levels with slightly less single peptide hits (**Figure 4C**).

### 4. LC-MS/MS method optimization

After sample collection and digestion conditions optimizations, we worked on chromatographic separation with the objective to reduce sample analysis time and thus increase sample throughput. To this end, we tested two separation column lengths: 25cm (75µm, 1.7µm, IonOptiks) with a 22-minute gradient, and 5 cm (75µm, 1.7µm, IonOptiks) with a reduced 10-minute gradient.

Comparative experiments showed that reducing gradient duration from 22 minutes (25 cm column) to 10 minutes (5 cm column) improved performance at lower injection amounts (Fig. 4). More precisely, with single cells replicates (N=10), an average of 3,782 protein groups were identified using the 5cm column, compared to 3,688 with the 25cm column. The shorter column configuration also demonstrates good consistency with a protein overlap of 85% (6,074 shared protein groups with the 25cm column). In addition, the coefficient of variation distributions are equivalent (Fig. 4C), confirming that the shorter column not only maintains sensitivity but also offers comparable quantitative performances at single cell equivalent quantities.

These optimizations, validated on single cell replicates, enabled a final throughput of up to 55 single cells per day (SPD), compared to 30 SPD with the 22 min gradient time configuration – with blanks and QCs included. While maintaining consistent coefficient of variation values ranging from 10% to 35%, this workflow offers in-depth coverage, reproducibility, and an improved analytical throughput, making it well-suited for large-scale single-cell proteomics studies. Moreover, this new chromatography configuration enables consistent high proteome coverage with no significant loss of performance, highlighting that the chromatographic retention miniaturization did not compromise data quality nor quantification accuracy.

## CONCLUSION

Through evaluation of sample preparation formats, we highlighted strengths and limitations of existing SCP workflows, including EVO96, LF48oil-based, and LF48oil-free. Then, the lysis and digestion step was evaluated and appeared to be crucial for final quantification results. Finally, nanoLC optimizations demonstrated that chromatography time can be reduced and thus the samples throughput improved with no significant loss of performance. All these optimizations led to the implementation of a sensitive and reproducible workflow, coupled with the nanoElute2, that addresses the main challenges of throughput, reproducibility, and sample handling in the SCP workflow (up to 5,000 protein groups were identified on HeLa cells at 55 SPD). We have also developed a sensitive, high throughput 120 SPD analysis workflow that can achieve up to 3,700 protein groups.

## ACKNOWLEDGEMENTS

The authors thank Dr. Christoph Krisp and Pierre-Olivier Schmit from Bruker Daltonics for their support and insightful advices on the instrumental settings. This work was supported by the Agence Nationale de la Recherche via the French Proteomic Infrastructure (ProFI UAR2048; ANR-10-INBS08-03 & ANR-24-INBS-0015), by the Region Grand-Est (SC-Proteomics project) and by the ITMO Cancer of Aviesan within the framework of the 2021-2030 Cancer Control Strategy, on funds administered by Inserm (ProteomiSC project), for equipment funds. It was also supported by the French Ministry of Higher Education and Research for the PhD fellowship of P.P.A and by the Interdisciplinary Thematic Institute IMS, the drug discovery and development institute, as part of the ITI 2021-2028 program of the University of Strasbourg, CNRS and Inserm, supported by IdEx Unistra (ANR-10-IDEX-0002), and by SFRI-STRAT’US project (ANR-20-SFRI-0012 for the PhD fellowship of C.K. C.C. is further supported by the European Union’s Horizon Europe MSCA PROHITS project under grant agreement (no. 101119980) and the CHIST-ERA project ODEEP-EU (G0GDV23N).

## SUPPLEMENTAL DATA

**Supplemental 1.**
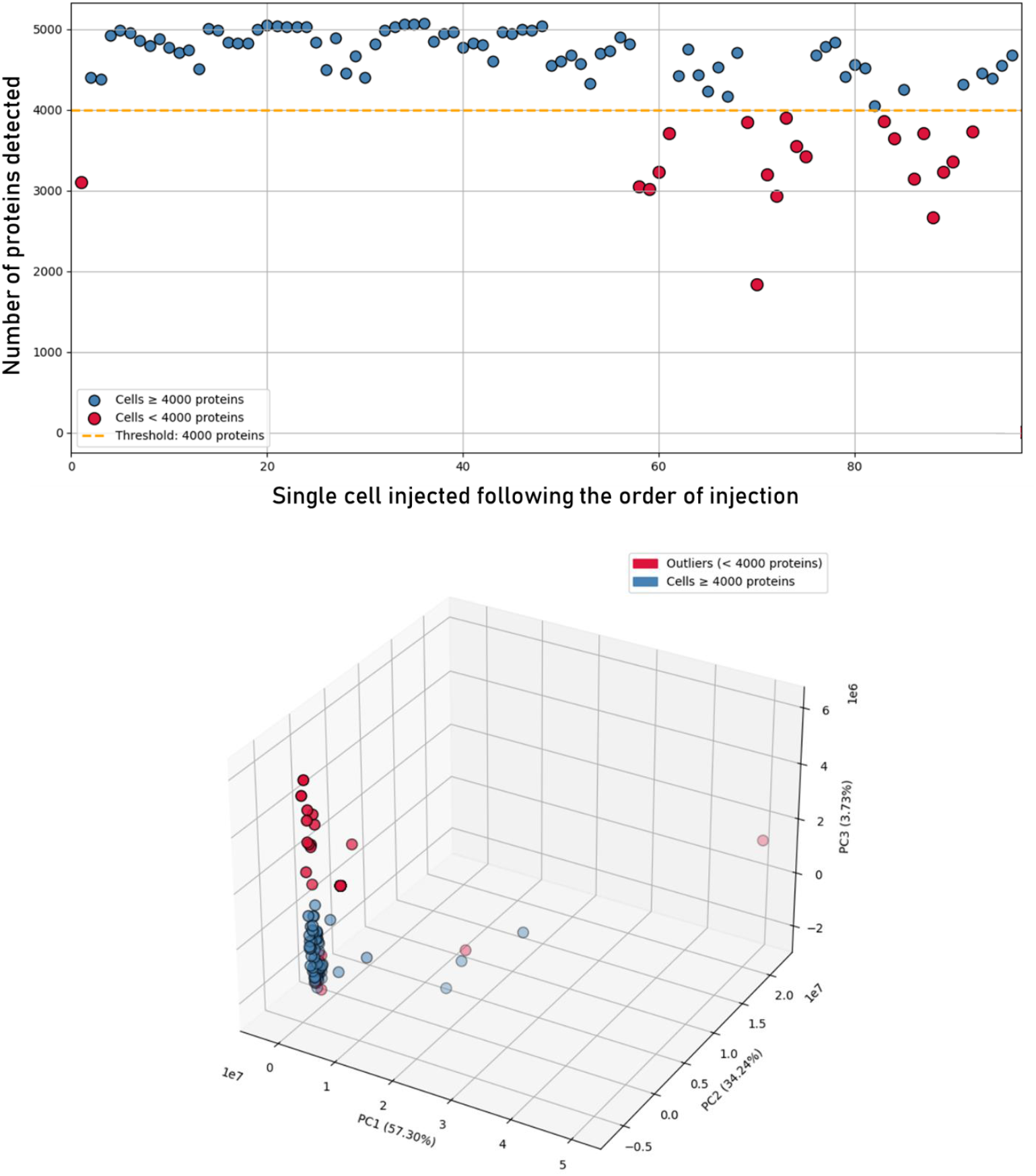
Single cell protein group numbers related to injection order for the oil-free LF48 direct injection workflow and Principal Component Analysis (PCA) of the 96 cells.

**Supplemental 2.**
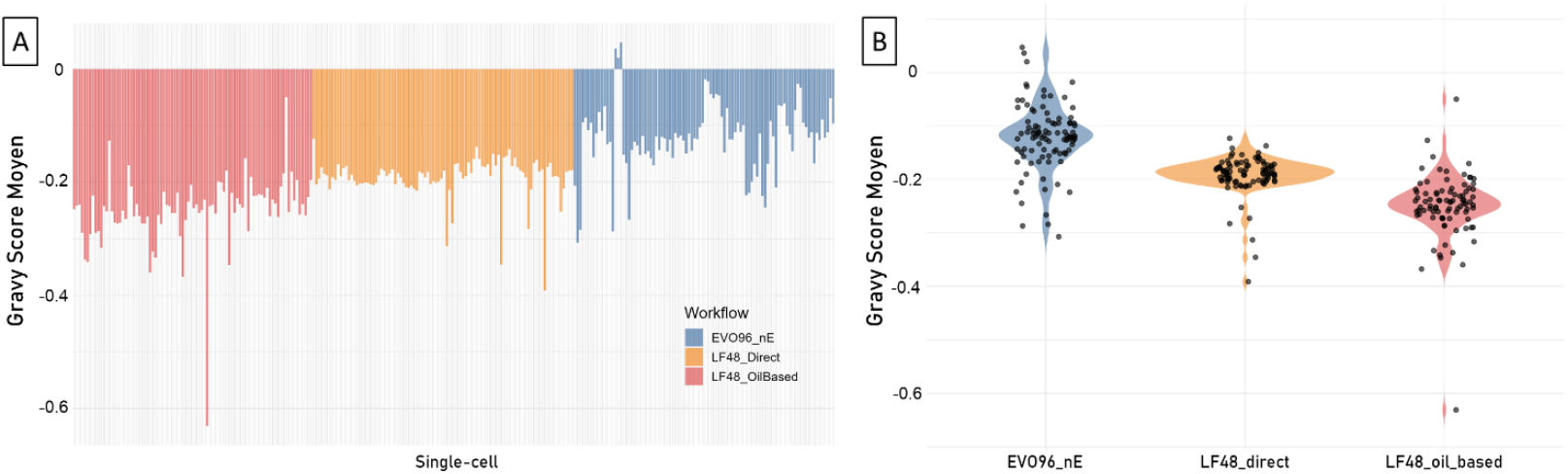
Repartition of GRAVY score across single-cells and workflows. To account for detection abundance, each peptide’s GRAVY score was weighted by its measured abundance in the corresponding cell, so that highly abundant peptides contribute more strongly to the cell’s mean GRAVY score, whereas low-abundance peptides contribute less. (A) Bar plot representing the distribution of the GRAVY scores across the three workflows. Each bar corresponds to an individual cells and its height indicates the weighted average GRAVY score of all peptides detected in that cell. (B) Violin plot showing the distribution of GRAVY scores across the three workflows.

**Supplemental 3.**
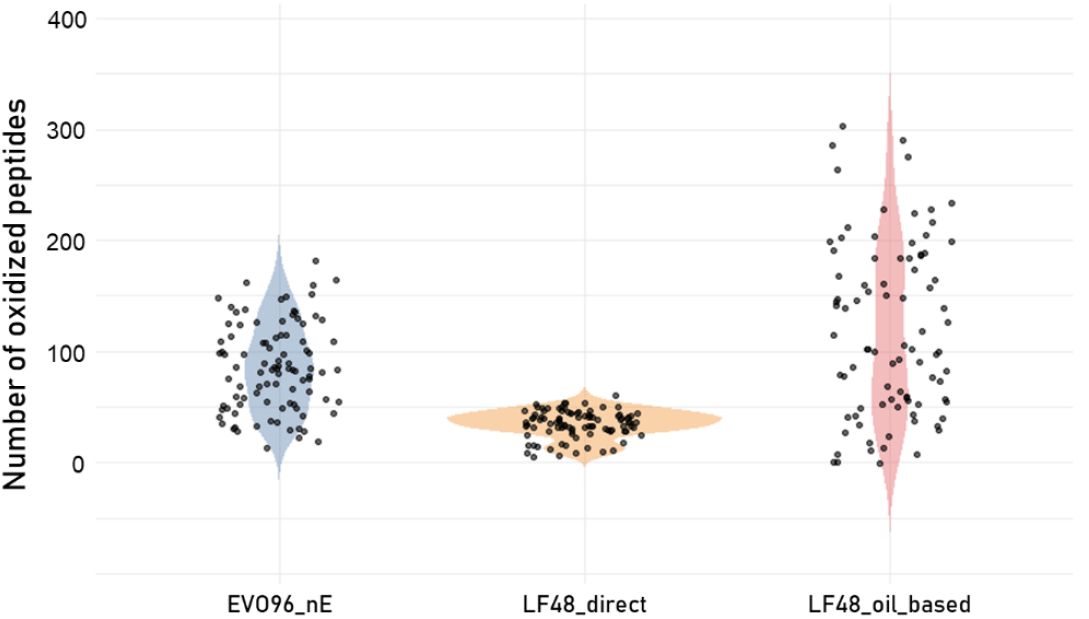
Distribution of methionine oxidations across workflows. Methionine oxidation phenomena were identified by searching the modified (UniMod:35) peptide sequences. Each dot represents the number of oxidized peptides detected in a single cell.

**Supplemental 4.**
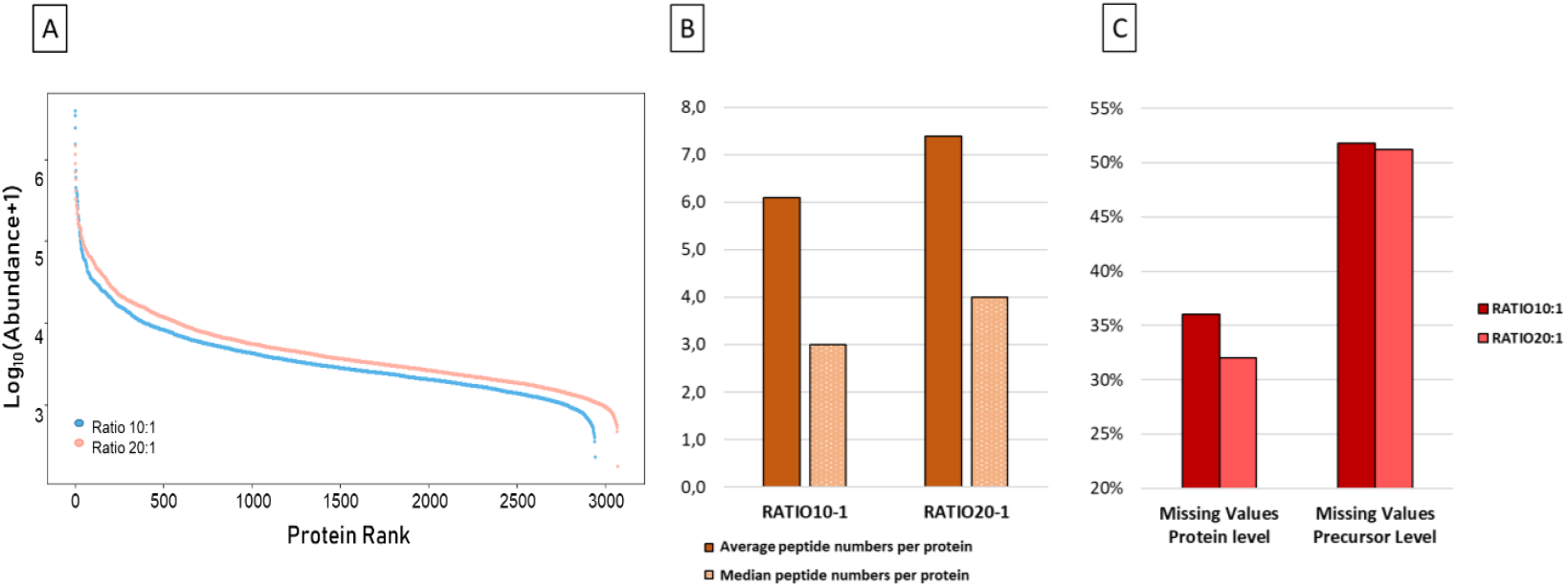
Impact of enzyme-to-protein ratios (10:1 and 20:1) on peptide and protein identifications on single HeLa cells (N= 96 per ratio). (A) Protein rank abundance distributions further compare the depth of proteome coverage under each digestion condition. (B) Average and median peptide numbers per protein for both ratios. (C) Missing values percentages at the protein and precursor levels for both ratios.

